# Changes in total charge on spike protein of SARS-CoV-2 in emerging lineages

**DOI:** 10.1101/2023.10.21.563433

**Authors:** Anže Božič, Rudolf Podgornik

## Abstract

**Motivation:** Charged amino acid residues on the spike protein of severe acute respiratory syndrome coronavirus 2 (SARS-CoV-2) have been shown to influence its binding to different cell surface receptors, its non-specific electrostatic interactions with the environment, and its structural stability and conformation. It is therefore important to obtain a good understanding of amino acid mutations that affect the total charge on the spike protein which have arisen across different SARS-CoV-2 lineages during the course of the virus’ evolution.

**Results:** We analyse the change in the number of ionizable amino acids and the corresponding total charge on the spike proteins of almost 2200 SARS-CoV-2 lineages that have emerged over the span of the pandemic. Our results show that the previously observed trend toward an increase in the positive charge on the spike protein of SARS-CoV-2 variants of concern has essentially stopped with the emergence of the early omicron variants. Furthermore, recently emerged lineages show a greater diversity in terms of their composition of ionizable amino acids. We also demonstrate that the patterns of change in the number of ionizable amino acids on the spike protein are characteristic of related lineages within the broader clade division of the SARS-CoV-2 phylogenetic tree. Due to the ubiquity of electrostatic interactions in the biological environment, our findings are relevant for a broad range of studies dealing with the structural stability of SARS-CoV-2 and its interactions with the environment.

**Availability:** The data underlying the article are available in the online Supplementary Material.

## 1. Introduction

The spike (S) protein of the severe acute respiratory syndrome coronavirus 2 (SARS-CoV-2) plays a role in target recognition, cellular entry, and endosomal escape of the virus (Huang et al., 2020). At the same time, the region in the viral genome that codes for the S protein is the one where most mutations tend to occur (Cavanagh, 2005; Harvey et al., 2021). Many of these mutations are intricately related to the changes in the structure of the S protein, thus influencing one or more of its functions (Mehra and Kepp, 2021). What is more, mutations on the S protein often lead to a change in its charge due to the ionizability of the corresponding amino acid (AA), and a series of recent studies has uncovered the tendency of these mutations to increase the total amount of positive charge on the S protein (Pawlowski, 2021, 2022; Cotten and Phan, 2023; Božič and Podgornik, 2023).

Changes in the amount of charge on the receptor-binding domain (RBD) of the S protein have been explored in detail, with a number of studies showing that they can lead to an increased binding of the virus to the angiotensin-converting enzyme 2 (ACE2) receptor (Adhikari et al., 2022a,b; Nie et al., 2022; Gan et al., 2022; Kim et al., 2023; Barroso da Silva et al., 2022). However, the *overall* distribution of charged AAs on the S protein is not trivial: Although the stalk part (S2 domain) is negatively charged, the top part of the spike molecule (S1 domain) and its RBD in particular remain predominantly positively charged for a broad range of pH values (Adamczyk et al., 2021; Kucherova et al., 2021). Surprisingly, the top of the spike reveals an underlying core with a surface area that has a dense negative charge (Kucherova et al., 2021), and the N-terminal domain (NTD) of the S1 domain also underwent several changes during the course of SARS-CoV-2 evolution making it net negative (Parsons and Acharya, 2023). Such a nonuniform charge distribution not only promotes spike corona stability, but also enhances virion attachment to receptors and membrane surfaces, which are mostly negatively charged (Adamczyk et al., 2021), and plays an important role in the non-specific electrostatic interactions of the virus with other charged macromolecules and macromolecular substrates in its environment (Javidpour et al., 2021; Arbeitman et al., 2021; Nie et al., 2021, 2022; Zhang et al., 2023).

That charge on the S protein is an important factor in the evolution of SARS-CoV-2 is further supported by a very interesting peculiarity, first identified by Paw-lowski (2021), who observed a tendency of the mutations in the S protein of different SARS-CoV-2 variants of concern (VOCs) to increase the number of positively charged AA residues. Subsequent studies by Cotten and Phan (2023) and Božič and Podgornik (2023) confirmed this finding on larger datasets; however, they also provided the first indications that this steady progression toward ever larger positive charge on the S protein was coming to a halt with the (then) newly emerging lineages. In order to resolve the question of whether the positive charge on the S protein of SARS-CoV-2 still has the capacity to increase as new lineages emerge, and to obtain a better understanding of the AA mutations that affect the total charge on the spike protein, we performed a study of the changes in the number of ionizable AAs on the S proteins of almost 2200 different SARS-CoV-2 lineages that have emerged between the start of the pandemic and end of December 2023. Our results show that not only has the steady increase in the amount of positive charge on the S protein come to a halt already with the early omicron variants, but that the pattern of AA changes and the resulting total charge have become more diverse with the newly emerged lineages. Clustering the lineages based solely on the patterns of change in the number of ionizable AAs on their S proteins shows that these patterns are highly characteristic of different major clades and make it possible to distinguish between broad clusters of lineages.

## 2. Materials and Methods

### 2.1 Data collection

Our data collection follows the approach described in our previous study (Božič and Podgornik, 2023): We first collected a list of SARS-CoV-2 Pango lineages from CoV-Lineages.org lineage report (O’Toole et al., 2021) on February 12th 2024. These were then used as an input to download SARS-CoV-2 genomic and protein data from NCBI Virus database (Hatcher et al., 2017) on February 13th 2024. Accompanying annotations were used to extract the isolate collection dates, where we kept the earliest *full* record (i.e., year, month, and day) as the timepoint of lineage emergence for use in our analysis. Lineage divergence— the number of mutations in the *entire genome* relative to the root of the phylogenetic tree, i.e., the start of the outbreak— was obtained from the global SARS-CoV-2 data available on Nextstrain.org (Hadfield et al., 2018). We selected only those entries with a genome coverage greater than 99 % and extracted their lineage divergence and the number of mutations. As the last step in the data selection, we retained only those Pango lineages whose downloaded fasta protein file was not empty. The total number of different SARS-CoV-2 lineages which passed this selection procedure and are included in our analysis is *N* = 2174. Since the number of S protein sequences included in the databases under each lineage varies, we consequently operate with average values taken over all sequences within a lineage; we denote these averages by 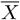

### 2.2 Ionizable AAs and their charge

We use Biopython (Cock et al., 2009) to parse the S protein fasta files and count the number of ionizable AAs on the S proteins of different SARS-CoV-2 lineages. We consider six different ionizable AA types: aspartic (ASP) and glutamic acid (GLU), tyrosine (TYR), arginine (ARG), lysine (LYS), and histidine (HIS). We omit cysteine (CYS), since it has a thiol with a functional end group that is a very weak acid and is typically not considered to be an acid at all (Nap et al., 2014; Božič and Podgornik, 2017).

Three of the six AA types carry positive charge (ARG, LYS, and HIS), while the other three carry negative charge (ASP, GLU, and TYR). Using their bulk dissociation constants (Table 1) and the Henderson-Hasselbalch equation (Božič and Podgornik, 2017), we obtain the charge on them at a given pH:

**Table 1.** Intrinsic pKa values of AA functional groups in bulk dilute aqueous solutions. Values taken from Haynes (2014).

|  | ASP | GLU | TYR | ARG | LYS | HIS |
| --- | --- | --- | --- | --- | --- | --- |
| $pK_a$ | 3.71 | 4.15 | 10.10 | 12.10 | 10.67 | 6.04 |

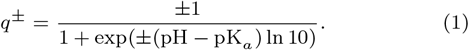

From Eq. (1) and Table 1, one can for instance discern that HIS typically carries a relatively small fractional charge at physiological pH, while TYR starts to acquire charge only at very basic pH. The total charge on the S protein is determined as the sum of the contributions of individual ionizable AAs,

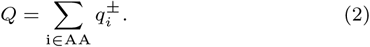

We are also interested in the relative change in the number of ionizable AAs on the S proteins of different lineages compared to the wild-type (WT) version of the S protein. To this purpose, we introduce the measure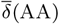,

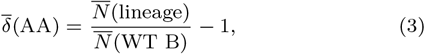

which quantifies the relative change in the number of each individual ionizable AA type on the S protein of a given lineage compared to their number on the S protein of WT lineage B.

## 3. Results

### 3.1 Changes in the number of ionizable AAs on the S protein of different lineages

We first take a look at how the number of different ionizable AAs on the S protein has been changing during the course of the virus’ evolution. Panel (a) of Fig. 1 shows the average number of ionizable AAs on the S protein as a function of the average lineage divergence, and thus also serves as a point of comparison to previous studies (Paw-lowski, 2021, 2022; Cotten and Phan, 2023; Božič and Podgornik, 2023). Our dataset includes SARS-CoV-2 lineages from the first appearance of the virus at the end of the year 2019 all the way to the lineages that have emerged as late as December 2023. Figure 1a already indicates that among the recently emerged lineages there exist several subpopulations, each of which prefers a specific number of ionizable AAs. For instance, the number of GLU seems to take on roughly four separate values for different lineages with divergence ≳ 70. We can also observe that the pattern of increase in the number of positively charged AAs is more prominent for LYS and HIS, and less so for ARG. Moreover, the change in the number of a given AA type can vary anywhere between 5 (e.g., GLU) to 10 (e.g., LYS) residues in the course of the evolution of different SARS-CoV-2 lineages.

**Fig. 1:**
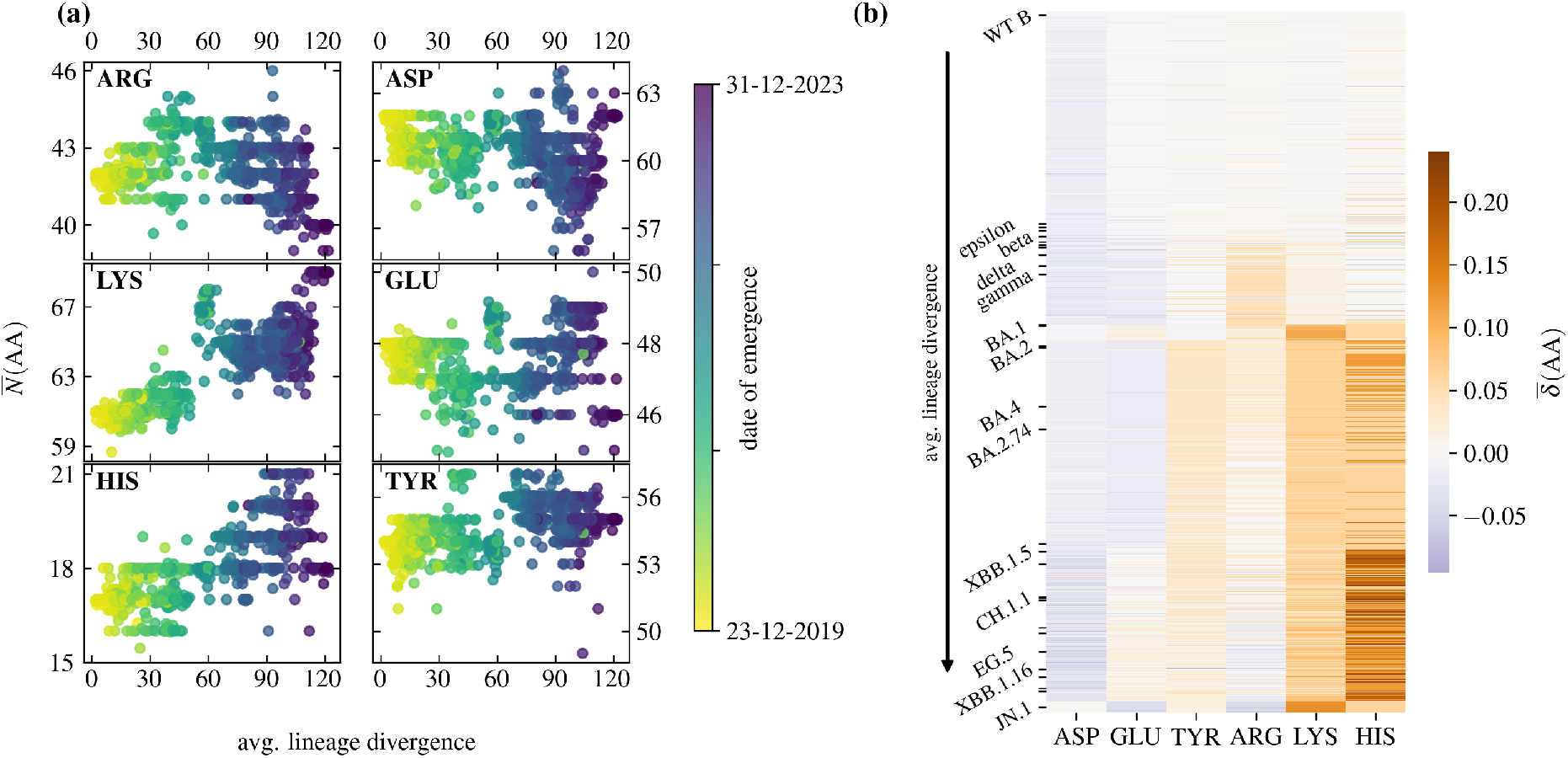
**(a)** Evolution of the number of ionizable AAs on the S protein with the divergence of SARS-CoV-2 lineages. AAs in the first column (ARG, LYS, and HIS) take on a positive charge, while AAs in the second column (ASP, GLU, and TYR) take on a negative charge. Each point represents the average value over the sequences from a single lineage, its colour corresponding to the first recorded date of its emergence. **(b)** Heatmap of the relative change in the number of ionizable AAs compared to the WT (lineage B), arranged according to lineage divergence [Eq. (3)]. Ticks along the y-axis mark select VOCs and VBMs, also listed in Table 2.

Different patterns of change in the number of ionizable AAs on the S protein are made more visible in the heatmap in panel (b) of Fig. 1, which shows the relative change in the number of ionizable AAs for each of the 2174 lineages analyzed when compared to the WT (lineage B) [Eq. (3)], arranged according to their average lineage divergence. Ticks on the *y* axis mark selected VOCs and variants being monitored (VBMs); these are listed in Table 2. While some changes seem prevalent across lineages, such as an increase in the number of LYS and TYR after the emergence of omicron, others—for instance, a significant increase in the number of HIS— do not occur regularly across all lineages, even though they are more likely to be seen in the recently emerged ones.

**Table 2.** Table of main current and previous SARS-CoV-2 VOCs and VBMs. Listed are the average divergence of the lineage (the average number of mutations with respect to the root of the phylogenetic tree), the earliest known date of emergence of the lineage, the average total charge on the S protein 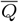, and the cluster to which the lineage belongs to based on the changes in the number of ionizable AAs compared to WT. Full table of all 2174 analysed lineages is available in the Supplementary Material.

| Lineage | Clade<sup>1</sup> | Avg. divergence<sup>2</sup> | Emergence<sup>3</sup> | $\bar{Q}$ [$e_0$]<sup>4</sup> | Cluster<sup>5</sup> |
| --- | --- | --- | --- | --- | --- |
| B | 19B | 4.29 | 2019-12-23 | -5.25 | 1 |
| B.1.427 | epsilon | 24.18 | 2020-09-17 | -3.28 | 1 |
| B.1.526 | iota | 25.22 | 2020-11-03 | -1.77 | 1 |
| P.2 | zeta | 25.74 | 2020-07-10 | -2.28 | 1 |
| B.1.351 | beta | 27.69 | 2020-10-09 | -1.41 | 1 |
| B.1.429 | epsilon | 28.07 | 2020-03-15 | -3.28 | 1 |
| B.1.525 | eta | 30.52 | 2020-12-15 | -1.30 | 2 |
| B.1.617.1 | kappa | 32.56 | 2021-02-08 | -0.39 | 2 |
| P.3 | theta | 33.03 | 2021-01-31 | -0.11 | 1 |
| C.37 | lambda | 33.80 | 2021-01-01 | -4.17 | 1 |
| B.1.1.529 | omicron | 36.74 | 2020-10-20 | -1.52 | 5 |
| B.1.621 | mu | 36.91 | 2021-04-20 | -1.36 | 1 |
| B.1.617.2 | delta | 38.73 | 2020-07-07 | -0.82 | 2 |
| P.1 | gamma | 39.58 | 2020-03-15 | -3.38 | 1 |
| BA.3 | 21M | 55.34 | 2021-12-04 | 2.82 | 5 |
| BA.1 | 21K | 56.12 | 2020-12-22 | 2.12 | 3 |
| BA.2 | 21L | 68.88 | 2021-12-08 | 1.85 | 4 |
| BA.5 | 22B | 69.06 | 2022-03-29 | 1.70 | 5 |
| BA.4 | 22A | 73.29 | 2020-09-02 | 1.78 | 5 |
| BA.2.74 | 21L | 74.88 | 2022-05-28 | 0.99 | 4 |
| BA.2.75 | 22D | 83.00 | 2022-01-04 | 1.76 | 6 |
| XBB | 22F | 86.05 | 2022-09-20 | 0.34 | 7 |
| XBF | recombinant | 90.08 | 2022-10-28 | 1.26 | 6 |
| CH.1.1 | 23C | 93.74 | 2022-09-29 | 1.16 | 6 |
| XBK | recombinant | 94.00 | 2022-11-07 | 1.22 | 6 |
| XBB.1.5 | 23A | 94.16 | 2022-11-14 | 0.33 | 7 |
| XBB.1.9.1 | 23D | 97.55 | 2022-12-29 | -0.22 | 7 |
| XBB.2.3 | 23E | 98.34 | 2022-12-21 | 1.04 | 7 |
| EG.5 | 23F | 101.28 | 2023-08-04 | 0.65 | 7 |
| XBB.1.16 | 23B | 104.17 | 2020-11-20 | 0.80 | 7 |
| FL.1.5.1 | 23D | 105.74 | 2022-10-12 | -0.08 | 7 |
| XBB.1.16.6 | 23B | 108.42 | 2023-05-07 | 1.92 | 7 |
| DV.7.1 | 23C | 108.98 | 2023-07-28 | 0.49 | 6 |
| BA.2.86 | 23I | 109.10 | 2023-08-03 | 1.55 | 8 |
| JN.1 | emerging | 118.79 | 2020-12-11 | 2.80 | 8 |

<sup>1</sup>Major variant or Nextclade (Aksamentov et al., 2021) clade designation.
<sup>2</sup>Average lineage divergence based on the number of mutations in the entire genome.
<sup>3</sup>Earliest full annotated date of lineage emergence.
<sup>4</sup>Average total charge on the S protein at pH = 7.
<sup>5</sup>Result of the $k$ -means clustering into 8 clusters based on relative changes in the number of ionizable AAs.

### 3.2. Patterns of change in the number of ionizable AAs

In order to better understand the different patterns of change in the number of ionizable AAs in different lineages seen in Fig. 1, we perform *k*-means clustering on the profiles of relative changes in the number of AAs, 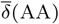, of all 2174 different lineages shown in Fig. 1b. In this way, we can observe which lineages are the most similar to each other in this respect. The optimal division of the lineages results in 8 clusters, and Fig. 2 shows their centroids— the mean relative changes in the number of ionizable AAs for all lineages contained in each cluster. Assignment of each lineage in our dataset to one of the eight clusters is available in Supplementary Material.

**Fig. 2:**
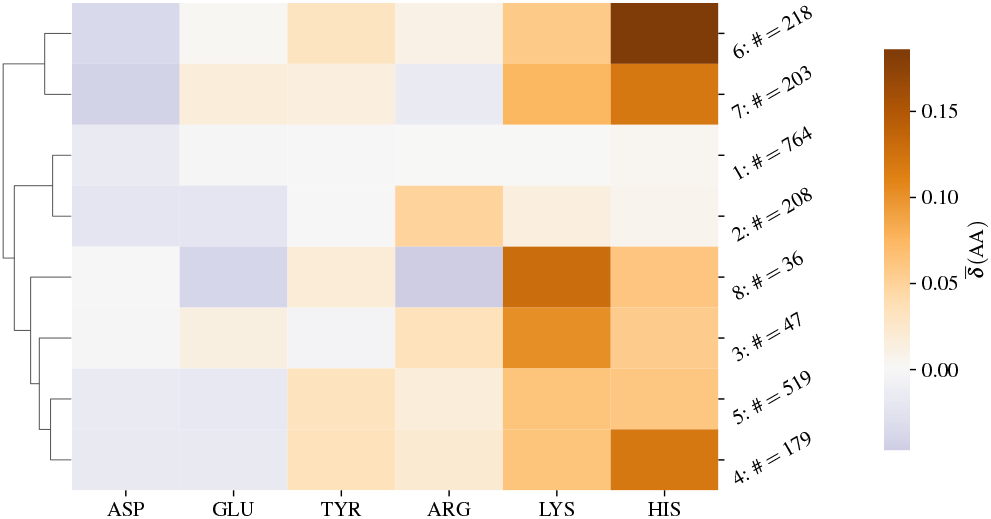
Centroids of the eight clusters obtained by *k*-means clustering of the relative changes in the number of ionizable AAs 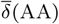, on the S proteins of 2174 SARS-CoV-2 lineages (Fig. 1b). ASP, GLU, and TYR take on a negative charge while ARG, LYS, and HIS take on a positive charge. Next to the numbered label of each cluster is the number of lineages in the dataset that belong to it. The centroids are further hierarchically ordered as shown by the dendrogram on the left side.

The largest cluster (cluster 1) shows almost no change in the number of ionizable AAs compared to the WT, while the cluster most similar to it, (cluster 2), already shows an increase in the number of (positively charged) ARG and a decrease in the number of (negatively charged) ASP and GLU—pointing towards an overall increase in the total charge on the S protein. This continues with cluster 3, where there is a significant increase in the number of LYS as well as an increase in the number of ARG and HIS. On the other hand, clusters 4 and 5, which are very similar to each other, show a decrease in the number of ASP and GLU, and an increase in the number of all other AA types. Clusters 6 and 7 are the most distinct from all the others, one exhibiting a larger increase in LYS compared to HIS whereas the other exhibiting the opposite. Finally, cluster 8 shows a decrease in both the number of GLU and ARG and the most significant increase in the number of LYS from among the eight clusters.

### 3.3. Total charge on the S protein of different lineages

Next, we wish to estimate how the patterns of change in the number of ionizable AAs on the S protein translate into the amount of charge that the S protein carries. To this purpose, we determine the total charge on the S protein for each lineage in a simple fashion by using the Henderson-Hasselbalch equation with bulk pK_*a*_ values for each AA type (i.e., without taking into account the specific environment of each residue—see also Materials and Methods and Discussion sections).

Figure 3 shows the average total charge on the S protein of 2174 different lineages as a function of their divergence for three different values of pH. Lineages are coloured according to the cluster they belong to (Fig. 2). Despite lineage divergence not being a parameter of the clustering, the first three clusters (1, 2, and 3) correspond well with the increase in lineage divergence. The S proteins of early lineages have an overall negative charge, and these lineages mostly fall under cluster 1 which shows fairly small changes in the number of ionizable AA compared to the WT (Fig. 2) and thus only a small increase in charge. The total charge nonetheless gradually increases with lineage divergence, and the spike proteins of lineages in cluster 2 already have a total charge around 0 at pH = 7. This cluster includes the early VOCs such as kappa and delta (Table 2 and Supplementary Material). Increasing the divergence still, the S proteins of lineages belonging to cluster 3 show quite an increase in charge, becoming overall positively charged at pH = 7; comparison with Fig. 2 indicates that this is likely mostly due to a large increase in the number of LYS. This cluster corresponds to the emergence of the early omicron lineages (Supplementary Material).

**Fig. 3:**
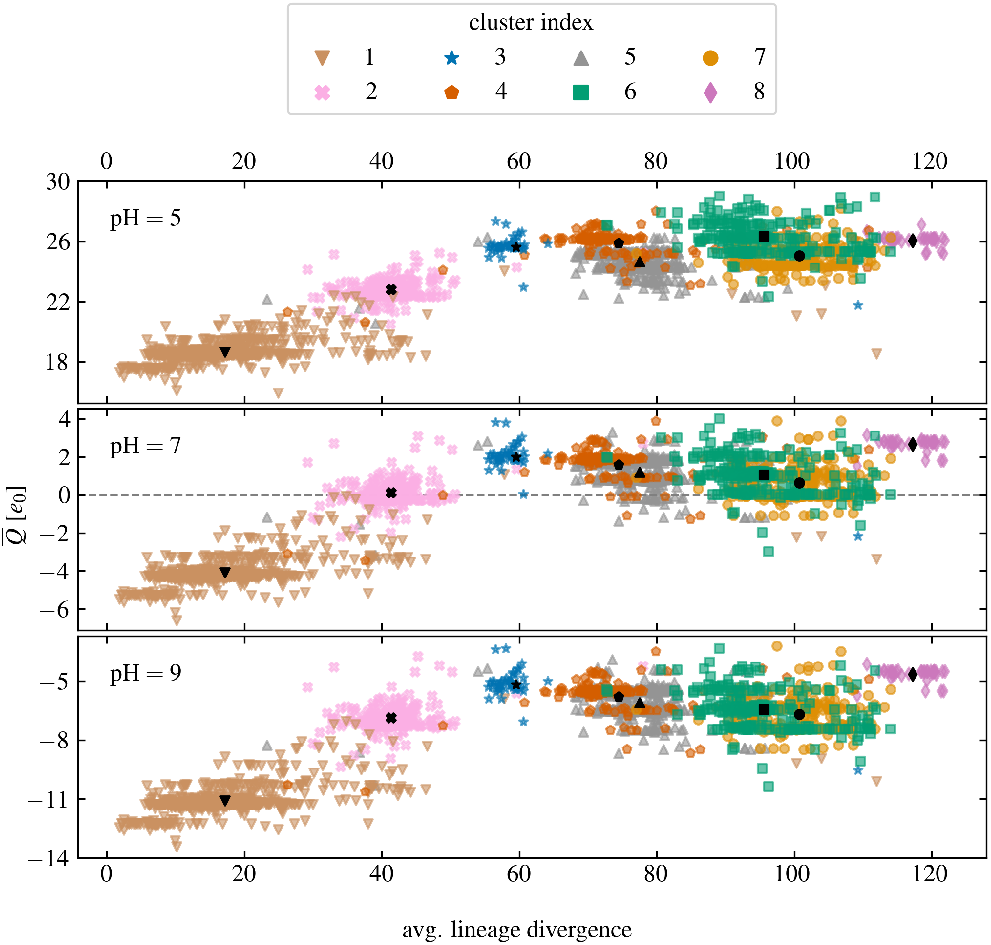
Average total charge on the S proteins of 2174 different SARS-CoV-2 lineages at three different values of pH (pH = 5, 7, 9) as a function of average lineage divergence. Individual lineages are coloured according to one of the eight clusters of their relative change in the number of ionizable AAs to which they belong (cf. Fig. 2). Black symbols show the total charge according to the centroid of each cluster and are positioned at the mean divergence of the cluster.

While the (average) changes in the number of ionizable AAs of clusters 4 and 5, which follow next, are different (Fig. 2), these changes do not lead to a significantly different amount of total charge on the S protein, albeit those belonging to cluster 5 have a slightly more positive charge. What is more, compared to the earlier cluster 3, the total charge on the S protein for the lineages in these two clusters is on average lower and shows a larger variability. Clusters 6 and 7 are similar in this respect: While their profiles of the average change of ionizable AAs are the most distinct from among all the clusters (Fig. 2), the total charge on the S protein they lead to is lower than the peak reached with lineages in cluster The total charge on the S proteins in these lineages is on average even lower than for the lineages in clusters 4 and 5. Similarly to these two clusters, the total charge on the S proteins of lineages in clusters 6 and 7 is also spread over a range of different values. Finally, the charge on the S proteins of cluster 8, which contains the most divergent lineages (including JN.1 and its subvariants), is again shifted towards more positive values, comparable to the charge seen in lineages in cluster 3. In this respect, lineages in cluster 8 represent a move away from the most recent trends in the changes in the number of ionizable AAs represented by lineages in clusters 6 and 7, with less variation in their charge and again a tendency towards an overall more positive charge.

### 3.4 Placing the changes in charge on the S protein into the phylogenetic tree of SARS-CoV-2 lineages

We can also place each of the analysed lineages on the general phylogenetic tree of SARS-CoV-2 sequences to investigate whether their division into clusters based on the changes in the number of ionizable AAs on their S proteins corresponds to the general phylogenetic division of their genomes. Figure 4 shows their placement on the phylogenetic tree generaged using Nextclade (Aksamentov et al., 2021), where each lineage is coloured according to the cluster it belongs to (Fig. 2). Interestingly, the clustering of the lineages—based *solely* on the changes in the number of ionizable AAs on the S proteins—corresponds extremely well to the broad division of lineages into different clades.

**Fig. 4:**
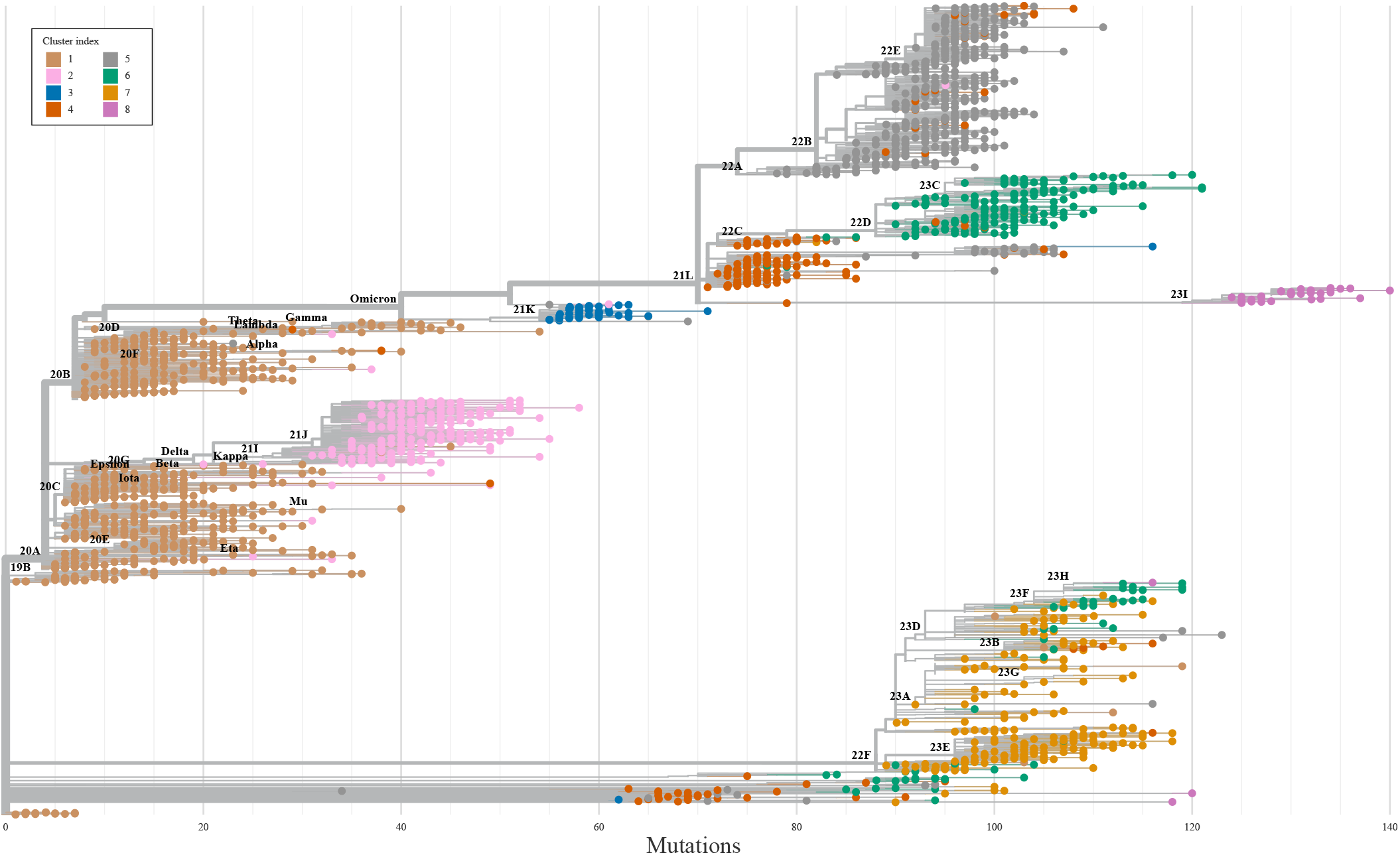
Placement of the 2174 SARS-CoV-2 lineages analysed in this work on the phylogenetic tree of all SARS-CoV-2 lineages as provided by Nextclade (Aksamentov et al., 2021). Only a single sequence of each analysed lineage is shown for clarity, and the tree is annotated with the major Nextstrain clades (Hadfield et al., 2018). Individual lineages are coloured according to one of the eight clusters of their relative change in the number of ionizable AAs to which they belong (cf. Fig. 2).

Cluster 1, with comparatively little change in the number of ionizable AAs compared to the WT, matches the earliest, least divergent clades of the phylogenetic tree, covering most VOCs up until omicron, as we have already noted before. Most of the lineages in cluster 2 belong to the major clade 21J and its descendants. Cluster 3 covers the clade 21K and indeed follows shortly after the emergence of omicron. Cluster 4 covers clades 22A, 22B, and 22E, while cluster 5 covers mainly the recombinant clade and a part of the clade 21L. Cluster 6 includes clades 22D and 23C, while cluster 7 covers recently emerged clades, including 22F, 23A (together with the so-called Kraken variant, XBB.1.5), 23B, 23D, 23E, and 23F. Cluster 8 includes clade 23I as well as the most recently diverged variant JN.1, yet unassigned to a clade.

Figures 3 and 4 together make it clear that while the total charge on the S protein of SARS-CoV-2 gradually became positive with the emergence of new lineages, it has reached a peak with the early omicron variants (cluster 3). At neutral and basic values of pH, more divergent lineages (clusters 4–8) retain the overall increase in the amount of positive charge on the S protein, but this charge has—with the exception of lineages in cluster 8—not only stopped increasing but also decreased slightly on average, as can be seen from the total charge according to the centroids of the clusters (black symbols in Fig. 3). In contrast, the mutations in the ionizable AAs on the S proteins of recently emerged lineages show that these mutations retain the increase in positive charge at acidic values of pH, even increasing it slightly further.

## 4. Discussion and conclusions

By studying the changes in the number of ionizable AAs on the S proteins of 2174 different SARS-CoV-2 lineages which have emerged between the start of the pandemic and end of December 2023 (Fig. 1), we have shown that these have initially led to a significant increase in the positive charge on the S protein which reached a peak with the emergence of early omicron variants. This trend has all but stopped with most newly emerged lineages—including the recent VBM Kraken (XBB.1.5)—in which the total charge of the S protein is still positive but shows a greater variability (Table 2 and Fig. 3). A notable exception to this is clade 23I, which includes the very recent VBM JN.1 and its subvariants. We have furthermore extracted 8 distinct patterns of relative change in the number of different ionizable AA types (Fig. 2), which not only closely follow the divergence of different lineages (Fig. 3) but also cover the broad division of major clades in the SARS-CoV-2 phylogenetic tree (Fig. 4). The latter observation is in line with previous studies which have demonstrated the possibility that VOCs could be identified from the sequence of the S protein alone, which would make lineage assignment relatively easier compared to the use of complete genome sequences (O’Toole et al., 2022).

Our work shows that the evolutionary changes in the ionizable AAs on the S protein have led to charge distributions which lead to different amount of total charge on the S protein as the solution pH shifts to acidic or basic values (Fig. 3). Given that it has been previously shown that the stability and activity of SARS-CoV-2 are significantly suppressed at very acidic and very basic pH values (pH ≲ 5 and pH ≳ 9, respectively) (Chan et al., 2020), it would be interesting to explore the functional importance of those mutations which retain high positive charge of the S protein at acidic pH. Such changes in pH can occur at different steps in the life cycle of the virus: For instance, the ability of the S protein’s fusion domain (FD) to mediate membrane fusion has a significant pH dependence, with fusion events more readily induced at low pH (*∼* 5–6) (Birtles et al., 2022; Kreutzberger et al., 2022). Initiation of fusion has also been found to be critically dependent on the protonation of HIS residues, which have in general been discussed as molecular switches due to their change from neutral to positively charged as the pH becomes acidic (Kampmann et al., 2006; Fritz et al., 2008; Qin et al., 2009). In contrast to FD, pH effects in S protein trimers at mild acidic pH have been found to be modulated mainly by ASP residues rather than HIS residues, where they serve to stabilize the locked form of the S protein at acidic pH (Lobo and Warwicker, 2021).

The total charge on the S protein that we determine here should be seen only as a proxy. While the values we find correspond to those obtained in studies using a similar approach (Pawl-owski, 2021, 2022; Cotten and Phan, 2023), studies using more detailed structural models that took into account local changes in the dissociation constants in general obtained higher values, even though these still vary among different studies (Adamczyk et al., 2021; Kim et al., 2023). Nonetheless, despite using a very simple model, we still observe the trends seen in studies which used more detailed models (Kim et al., 2023; Barroso da Silva et al., 2022). Thus, while the values of charge on the S protein we determine are certainly not exact, the patterns we see should translate even to studies using more accurate models for the charge on the AA residues. To truly determine the charge on each individual AA residue would require not only the knowledge of the the correct structure of the S protein for each and every lineage (Mehra and Kepp, 2021; Casalino et al., 2020), which is impossible to obtain, but one would also need to resort to simulations based on density functional theory (Ching et al., 2022; Jawad et al., 2022; Ching et al., 2023), which can be computationally prohibitive even for a single S protein structure, let alone for such a large dataset.

Effects of specific mutations and related changes in local charge also require a detailed structural analysis and are thus beyond the scope of our work. However, we can utilize the clustering of changes in ionizable AAs (Fig. 2 and its correspondence with the major clades of SARS-CoV-2, Fig. 4) to connect these changes with their approximate location on the S protein, as illustrated in Fig. 5. Panels (a)–(d) show the electrostatic potential on the S protein of four select SARS-CoV-2 VOCs, where we can observe both the trend towards higher positive charge on the S proteins of variants which have emerged later as well as the fact that the mutations which give rise to this trend are not localized to a specific region but can be found on different parts of the S protein structure. We can further explore this by relating the positions of *characteristic* mutations in ionizable AAs to the main structural domains of the S protein, shown in panel (e) of Fig 5. Mutations which either change an ionizable AA to a neutral/ionizable one or vice versa are shown for eight variants belonging to each of the clusters we identified in this work (Fig. 2). Most changes in the number of ionizable AAs can be seen to occur in the S1 domain of the S protein—predominantly in the NTD and RBD regions—and only a few of them occur in the S2 domain. Moreover, the mutations in the NTD region tend to make the charge more negative (Parsons and Acharya, 2023), whereas those in the RBD region make it more positive, although opposite changes can be observed as well. It is therefore all that more interesting that despite the occurrence of both mutations which increase and those which decrease the local charge, the overall charge on the S protein has been evolving towards progressively higher values during the course of the pandemic.

**Fig. 5:**
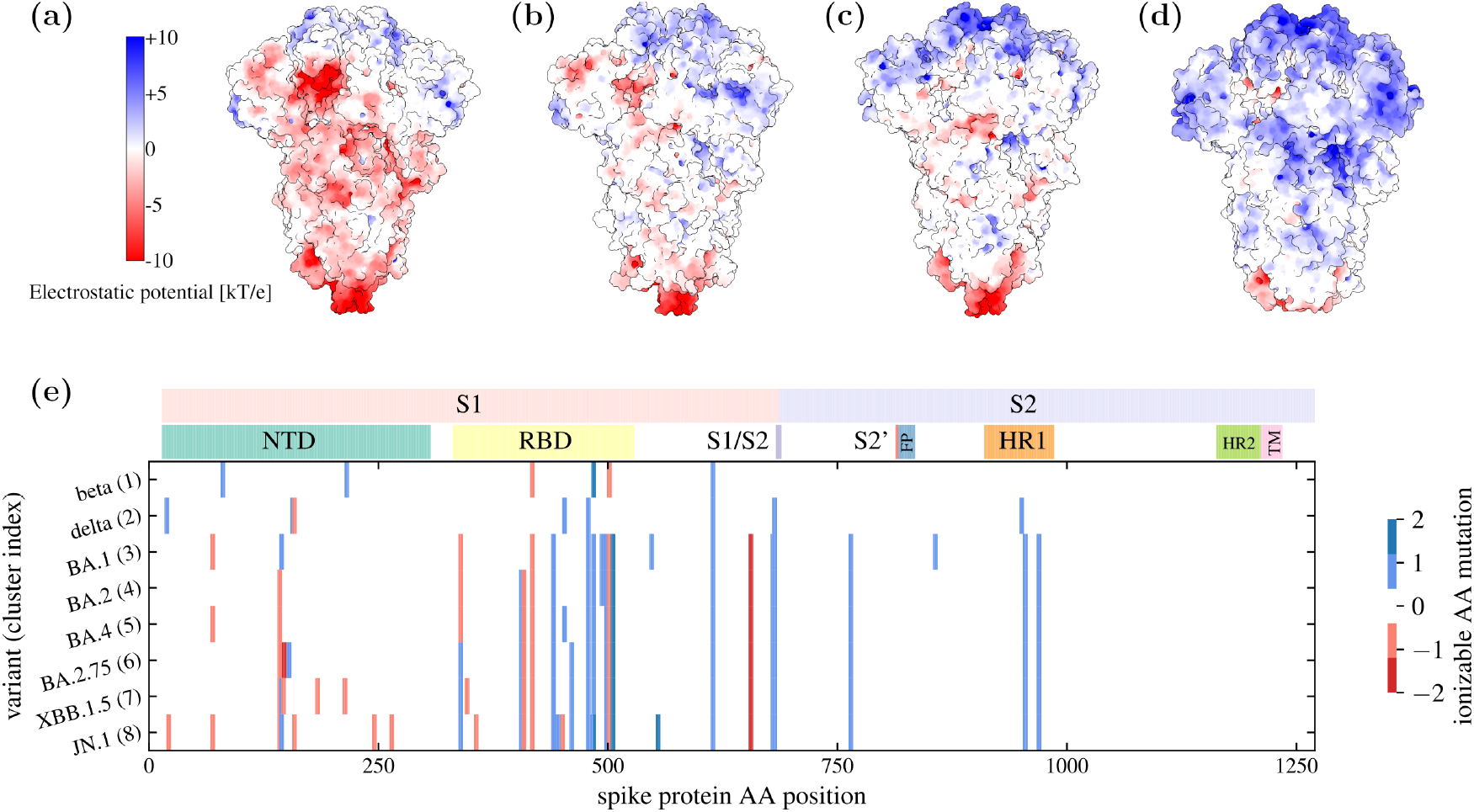
**(a)**–**(d)** Electrostatic surface potential on the S proteins of four different SARS-CoV-2 variants: (a) WT [PDB: 7FB0], delta [PDB: 7TUO], omicron BA.2 [PDB: 7UB6], and omicron BA.2.75 [PDB: 8GS6]. The electrostatic potential was determined at pH = 7 using APBS and PDB2PQR software (Jurrus et al., 2018). **(e)** Characteristic mutations of ionizable AAs on the S proteins of different SARS-CoV-2 variants, each variant belonging to one of the eight charge clusters identified in this work (Fig. 2; cluster index is noted in parentheses next to the variant name). The variant-characteristic mutations were obtained from cov-spectrum.org, and only those mutations occurring in at least 70% of the sequences were retained. Mutations are marked as +1 if the change is from a negatively charged AA to a neutral one or from a neutral one to a positively charged one; as *−*1 if the charge is from a positively charged AA to a neutral one or from a neutral one to a negatively charged one; and as *±*2 if the change is from a negatively charged AA to a positively charged one or vice versa. The positions of the mutations on the S protein AA sequence are annotated with some of the most important regions of the S protein (Jackson et al., 2022): NTD – N-terminal domain; RBD – receptor-binding domain; S1/S2 – furin cleavage site; S2’ – S2’ cleavage site; FP – fusion peptide and fusion-peptide proximal region; HR – hexad repeat; TM – transmembrane anchor.

Finally, it is worth mentioning that mutations contributing to the positive charge of SARS-CoV-2 have been recently identified as particularly important in the context of host antimicrobial peptides (xenoAMPs), composed of SARS-CoV-2 protein fragments that can co-assemble with double-stranded RNA into a form of proinflammatory nanocrystals, capable of multivalent immune recognition and a drastically amplified inflammatory response (Zhang et al., 2024). This particular consequence of changes in the charge of SARS-CoV-2 would imply that, apart from the indisputable role of individual mutations in the specific interactions between the S protein and the ACE2 receptor, the *overall* charge on the S proteins can also play a comparably important role to indicate whether a coronavirus can lead to a pandemic and cause severe inflammation, or simply result in symptoms akin to a common cold (Gussow et al., 2020; Božič and Podgornik, 2023). It could be then that this aspect of the cationic amplification in the evolution of SARS-CoV-2 S protein plays an even more important role than the modifications wrought in the specific interactions between the virus and cellular receptors.

The results of our work are important for a broad range of studies involving SARS-CoV-2: The relevance of the changes in the number of ionizable AAs on the S protein goes beyond those which occur in the RBD of the protein and which have direct implications for its binding to the ACE2 receptor. Some of these mutations can play a role in (de)stabilizing or altering the structure of the S protein in either closed or open states (Warwicker, 2021; Boer et al., 2022), while mutations occurring in other parts of the S protein can be a consequence of adaptation to more general electrostatic interactions with the environment of the virus, for instance with lipid rafts in cellular membranes (Matveeva et al., 2023). Lastly, overall accumulation of positive charge in SARS-CoV-2 and other coronaviruses as such has been linked to high fatality rates (Gussow et al., 2020) and can be an important factor in the assembly of xenoAMP proinflammatory nanocrystals which include SARS-CoV-2 protein fragments (Zhang et al., 2024).

## Data availability

Data presented in this work are available in the online Supplementary Material.

## Author contributions statement

R.P. conceived the study and A.B. performed the analysis and analysed the results. A.B. and R.P. wrote and reviewed the manuscript.

## Acknowledgments

We thank Domen Vaupotič for help with preparing Figure 4.

